# Cephalo-pelvic covariation and sexual dimorphism are disrupted in hybrid mice: implications for the human obstetrical dilemma

**DOI:** 10.64898/2026.05.11.724362

**Authors:** Eva Zaffarini, Kerryn Warren, Marta Vidal-García, Rebecca R Ackermann, Barbara Fischer, Philipp Mitteroecker, Benedikt Hallgrímsson

## Abstract

Cephalo-pelvic disproportion in humans has traditionally been interpreted through the obstetrical dilemma framework, assuming a trade-off between bipedal locomotion and childbirth. However, cephalo-pelvic covariation and pelvic sexual dimorphism might be common adaptations to parturition among mammals. We use a controlled hybridization model in mice to test whether cephalo-pelvic covariation and pelvic sexual dimorphism are population-specific, genetically structured, and sensitive to hybridization. We analyzed skull-pelvis variation and covariation, as well as sexual dimorphism of pelvic morphology across four divergent wild-derived mouse strains and their hybrids. Hybridization induced consistent cranial and pelvic size enlargement. Females exhibited significant cephalo-pelvic shape covariation, characterized by an association between rounder, wider birth canals and larger neurocrania, consistent with functional integration under obstetric selection. Hybrids showed disrupted size covariation, increased pelvis shape variance, and reduced cephalo-pelvic integration. Pelvic sexual dimorphism was systematically reduced in hybrids. Cephalo-pelvic covariation and pelvic sexual dimorphism are not exclusive to bipedal or encephalized species. They likely reflect widespread selection on birth canal morphology in mammals and have a complex genetic basis sensitive to hybridization. These findings weaken a human-exclusive interpretation of the obstetrical dilemma and highlight genetic introgression as an understudied factor shaping cephalo-pelvic integration and disproportion risk in mammals, including humans.

## Introduction

Cephalo-pelvic disproportion, a mismatch between the neonate’s head and the maternal birth canal, is the leading cause of obstructed labour worldwide (1). Evolutionary causes of cephalo-pelvic disproportion have been attributed to the “obstetrical dilemma” hypothesis: human birth canals partially trade off an increase in size to facilitate the birth of large babies with narrow pelvises for efficient bipedal walking and pelvic floor stability. The resulting pelvis is considered obstetrically insufficient and at risk of obstructed labour (2,3).

The risk of cephalo-pelvic disproportion is evolutionarily alleviated in humans by two complementary strategies. Firstly, the covariation of maternal pelvic canal and neonatal head morphology reduce the negative impact of the obstetrical dilemma and the risk of cephalo-pelvic disproportion. Women with relatively larger cranial dimensions tend to have a proportionally wider pelvic canal, especially in the pelvic outlet, facilitating the delivery of neonates with similarly large head size (4). The covariation of head and pelvic morphology is subject to selection to reduce the negative consequences of morphological mismatch (5). Secondly females show a rounder, slightly wider pelvic canal compared to males, a shape selected to facilitates the descent and rotation of the neonate during parturition (3,4,6,7).

Covariation among traits with related functions is a well-documented evolutionary phenomenon (8). Cheverud (1996) argued that selection on the relative proportions of traits that share a common function tends to promote the selection of pleiotropic genes and developmental processes (9). Therefore, selection for skull and pelvis covariance between mother and fetus and within adult individuals should have promoted genetic integration between these two traits in humans. Recently, the genetic basis for covariation between head shape and pelvis shape has been revealed in humans. In particular, covariation of adult head width and pelvic inlet width corresponds to an equally significant covariation with fetal birth weight, used as a proxy for fetal head size (10).

The extent of cephalo-pelvic morphological and genetic covariation might vary in different populations. Reported studies found high morphological covariation in a North American sample (4), significant genetic correlation in a UK sample (10) and non-significant morphological covariation for a Brazilian sample (11). Pelvis shape, head shape and fetal size are influenced by genetic background, ecological and life history variables (12–15). This means that different populations are expected to evolve distinct, genetically structured patterns of morphological integration between head shape and pelvis shape. Pelvic sexual dimorphism, likewise, shows a complex genetic architecture, is geographically structured, and shows variations in patterns following environmental and sexual selection as well as neutral genetic variation (10,16–19).

Pelvic sexual dimorphism and cephalo-pelvic covariation are expected to evolve in populations of mammals with a tight cephalo-pelvic fit (5,20–22). Small-bodied primates, like squirrel monkeys, bushbabies and macaques, display a high feto-maternal size ratio and strong covariation between neonatal head shape and maternal birth canal shape (23,24). In a simulated parturition model using the true maximum diameter of the chimpanzee birth canal, chimpanzee fetal heads negotiate birth canals previously considered wider, suggesting a certain degree of feto-pelvic covariance could be selected positively (25).

However, it is unclear whether pelvic sexual dimorphism evolves exclusively in response to obstetric selection. Pelvis sexual dimorphism has been found in primates with more difficult childbirth (hylobates, macaques) (14,25–30), as well as chimpanzees which give birth to relatively small neonates (31). Even, marsupials like kangaroos, koalas and wallabies, which give birth to extremely small and altricial neonates, show pronounced sex differences in pelvic morphology (26,32). In rodents, independently of the size of their litter, the already present pelvic sexual dimorphism is further exacerbated by pregnancy (33). Sex differences in these mammals might be justified as either retention of an ancestral dimorphism in absence of selective pressures, or as the product of mild but significant obstetric selection (25,31,34)

Since pelvis shape and skull shape vary significantly among populations, variance and covariance in these traits might be genetically and geographically structured. However, contrary to humans, very little is known about the genetic and developmental basis of cephalo-pelvic covariance and sexual dimorphism in non-human mammals.

We used a mouse model of hybridization to test whether cephalo-pelvic covariation is structured in population-specific ways. Hybridization among distantly related populations can cause perturbations of the underlying genetics and variance of the processes that drive covariation among functionally related traits (35–38). This is true for cephalo-pelvic disproportion as studies of domestic animal hybrids that show elevated risk for birth complications (39–42). If the developmental-genetic mechanisms underlying increased size are different in founder strains, hybridization may increase the variance of traits in ways unrelated to function. This can be complicated by heterosis (higher values for hybrid trait values compared to parental values) in which the offspring exhibits disproportionate body size (43,44). Hybridization is also known to affect sexual dimorphism, by either increasing or decreasing it (45), affecting the risk of obstructed labour independently of cephalo-pelvic covariation.

Mice of the genus *Mus* are an attractive model for the study of cephalo-pelvic disproportion due to extensive knowledge of mouse genetics and the existence of systematic crosses among distantly related strains. Like humans and other mammals, mice experience obstructive dystocia, often due to large fetus (46). However, their quadrupedal mode of locomotion is unlikely to generate the same degree of selective trade-off between locomotor efficiency and pelvic shape as has been proposed for humans. Mice also show pelvic sexual dimorphism, exacerbated by pregnancy, large litters, and craniometric size (33,47). All these factors suggest that mice are also selected for obstetrically efficient birth canal morphology.

We take advantage of four divergent wild derived strains of mice (*Mus mus musculus, M. m. castaneus, M. m. domesticus and M. spretus*) that vary significantly in body size, craniofacial and pelvic morphology. We examine the effect of hybridization on cephalo-pelvic covariation in relation to the functional demands of parturition and on the degree and pattern of sexual dimorphism. Specifically, we test the following hypotheses: i) Mice present skull-pelvis covariance consistent with selection for obstetrical demands, specifically in females. We predict large skulls will be associated with rounder, wider birth canals; ii) the covariance of skull and pelvis shape is the highest in F0 and reduced in hybrid populations. If the determinants of covariation between head and pelvis within each strain have diverged, then the hybrids should exhibit reduced covariation; iii) mice present pelvic sexual dimorphism consistent with selection for obstetric needs. We predict that female pelvises will present a series of pelvic features consistent with the overall enlargement of the birth canal compared to male; iv) pelvic sexual dimorphism is altered in hybrids in both magnitude and pattern, compared to F0.

## Results

### Hybrid neurocranial size increase is not matched by pelvis size increase

We first measured pelvis and neurocranium size for one generation of parental strains (F0), two hybrid generations (F1, F2) and three backcrosses (B1, B2, B3). Hybrids and backcrosses exhibited consistently larger pelvis and neurocranium sizes compared to F0 (Fig. 1). Pelvic centroid size increased on average by 2.54% (0.91% in females; 4.00% in males), whereas neurocranial centroid size increased by 1.21% (1.09% in females; 1.32% in males), indicating a stronger size response in the pelvis, particularly in males. Generation-specific patterns of pelvic and neurocranial size highlight heterogeneity between sexes and generations. F1 hybrids showed nearly identical proportional increases in pelvic and neurocranial size in both sexes. In females, F2 and B1 displayed comparable size increases in both traits, whereas B2 and B3 exhibited a marked enlargement of the neurocranium with little or no corresponding change in pelvic size, resulting in a decoupling of cranial and pelvic growth. In females, F0 pelvic size differed significantly only from F1, whereas neurocranial size differed significantly from all generations except F2.

**Fig 1.**
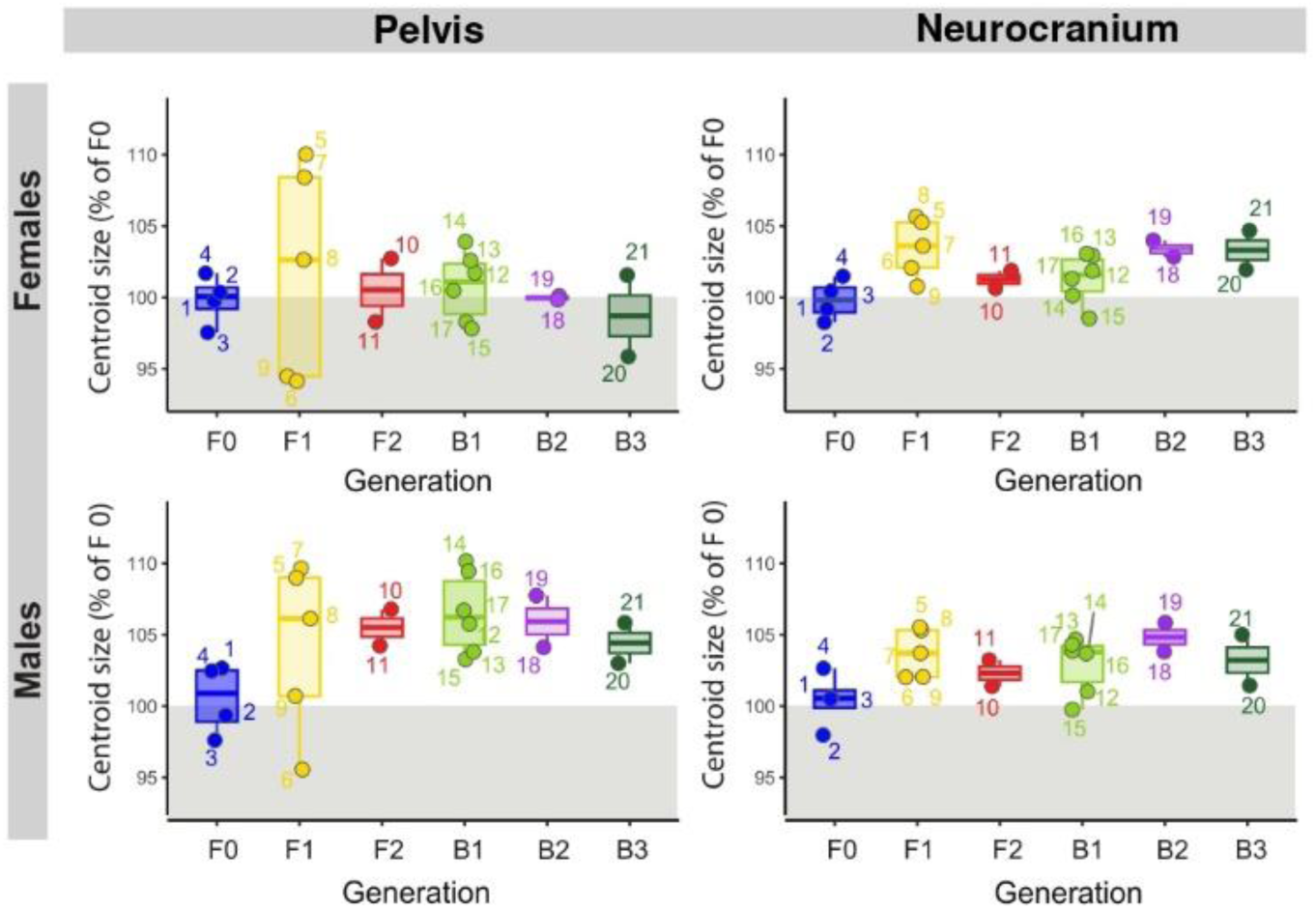
Mean pelvis and neurocranium centroid size for each generation. Neurocranium size (right column) increases at every generation for both sexes. Pelvis size (left column) increases only in males, while female hybrids tend to have a centroid size similar to F0 females. The values are expressed as percentage of the mean centroid size of F0. Each dot represents the mean centroid size for each strain. Significant differences between generations are reported in Supplementary Figure 4 as asterisks (p<0.05*, p<0.01**, p<0.001***).

### Pelvic and cranial morphologies respond differently to hybridization

Principal Components Analysis (PCA) of cranial and pelvis shape, standardized by mean generation shape, revealed distinct variance structures between traits and between sexes. PC1 of pelvis shape (36.73%, Fig 2a, left column), captured the variation between pelvises with long sacrum, closed pubic symphysis, thick pubic bones and small birth canal (Fig. 2b), and pelvises with short sacrum, thin and caudally extended pubic bones, open pubic symphysis, short ilia, and a wide birth canal. PC2 (12.42% of pelvis shape variance) highlighted a widening of the ischia and an open pubic symphysis, and a caudally tilted sacrum on one extreme of the axis and pelvises with narrower ischia, closed pubic symphysis and ventrally tilted sacrum on the other extreme. When the PCA was repeated after correcting for strain mean shape instead of generation (Supp Fig. 6), the variance structure became more similar between skull and pelvis. Under this normalization, F2 individuals of both sexes consistently showed the greatest dispersion along the first two PCs for both traits, indicating elevated shape variance compared to F0 and all other generations (Supplementary Fig 6).

**Fig. 2.**
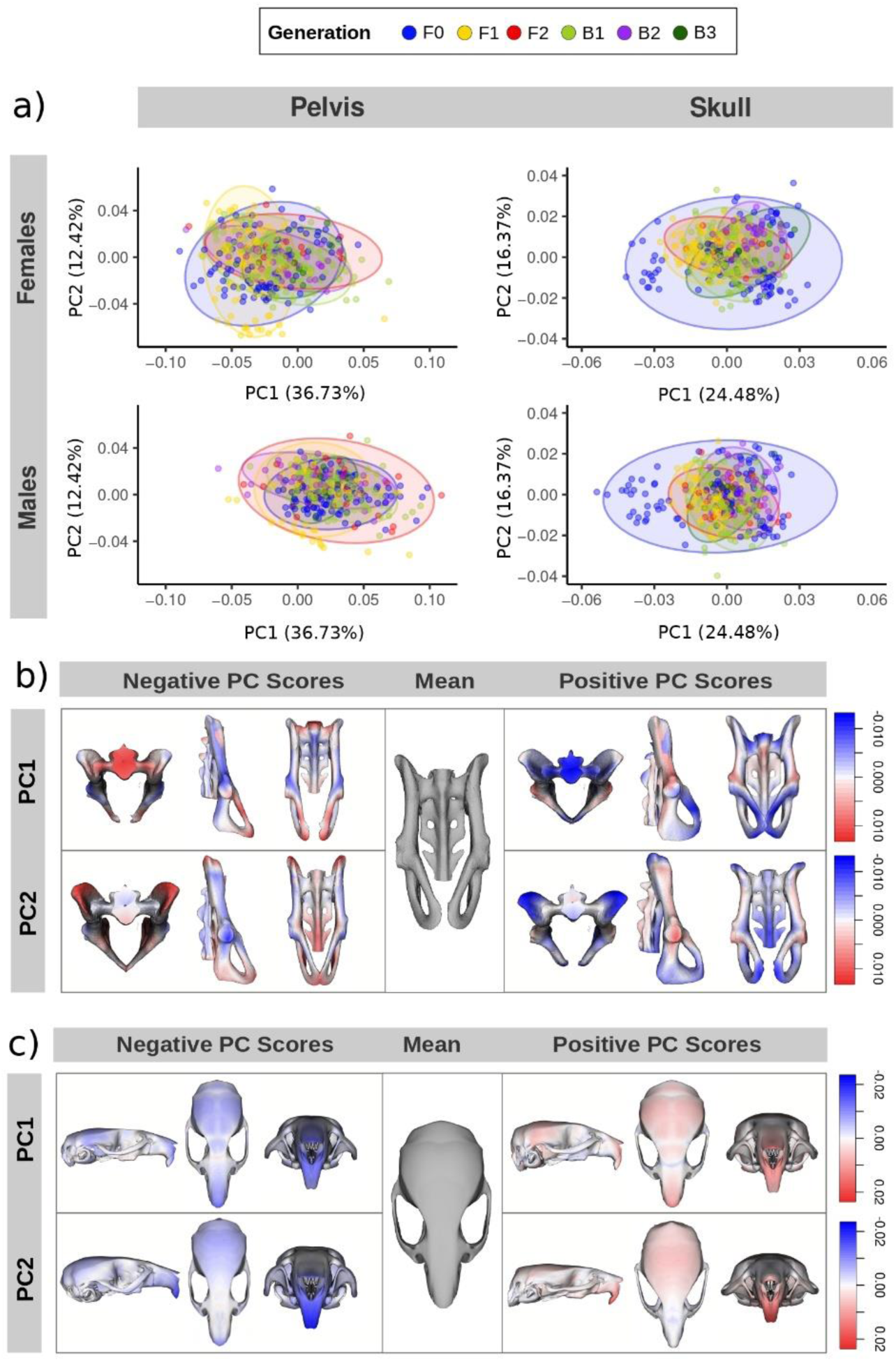
Principal component analysis of pelvis and skull shape. Across generations (panel a), hybrids display more extreme pelvic phenotypes than F0 (blue ellipses), whereas F0 shows the greatest cranial shape variance. (*a*) PCA of pelvis and skull shape for females (top), and males (bottom), mean-centered for generation. Colored ellipses denote 95% confidence intervals of individual distributions. (*b*) Mean pelvis shape (grey) with exaggerated morphs along PC1 and PC2 shown in transversal, sagittal, and coronal views, based on the generation-centered PCA. Morphs were generated by scaling PC loading vectors by 4× their respective eigenvalues. (*c*) Mean skull shape (grey) with exaggerated morphs along PC1 and PC2 in the same views. Morphs were scaled by 5× (PC1) and 10× (PC2) eigenvalues. Heatmaps indicate expansion (red) and contraction (blue) relative to the mean.

Extreme phenotypes emerged in both sexes. Female F1 hybrids had more extreme positive values of PC2, corresponding to the appearance of wider birth canals with an open pubic symphysis, whereas F2 and B1 females exhibited extreme positive values along PC1, relative to F0 (Fig. 2a) corresponding to closed pubic symphysis and elongated sacrum (Fig 2b).

Cranial shape showed a markedly different pattern. For the skull, PC1 and PC2 accounted for 24.48% and 16.37% of total variance, after correction for mean generation shape (Fig 2, right). F0 individuals spanned most of the variation along PC1 and PC2, whereas hybrids clustered near mean PC values. PC1 shows cranial shape variation mostly affecting the length and width of the skull (elongated and narrow versus shortened and wider), whereas shape variation along PC2 reflects a “shearing” effect on the whole skull (Fig. 2c). When correcting for mean strain shape instead of mean generation shape, F2 crania showed the highest level of dispersion and presence of extreme phenotypes compared to the other generations, including F0 (Fig 2, right and Supp. Fig. 6).

### Female mice show substantial covariation between pelvis and skull morphology

In females, the covariance of skull and pelvis analysed through Partial Least Square Analysis (PLS) was significant (*r*=0.34, *p*=0.001), and the first PLS axis explained 32% of the total skull-pelvis covariation (Fig. 3). After correlating the scores for skull shapes onto scores for pelvis shape, we found that F0 and B1 showed significant correlations, while F1, F2, B2, and B3 showed non-significant correlations. The Mean Square Error (MSE, a measure of dispersion around the covariance axis) was highest in F2 and lowest in F1 (Supp. Table 14). Pelvises with a long sacrum, narrow birth canal and small pubic gap were associated with flattened neurocrania, long face relative to the neurocranium, and prominent cheekbones. Pelvises with a wide and round birth canal, open pubic symphysis and dorsally positioned sacrum were associated to skulls with a wide neurocranium and short face.

**Fig. 3.**
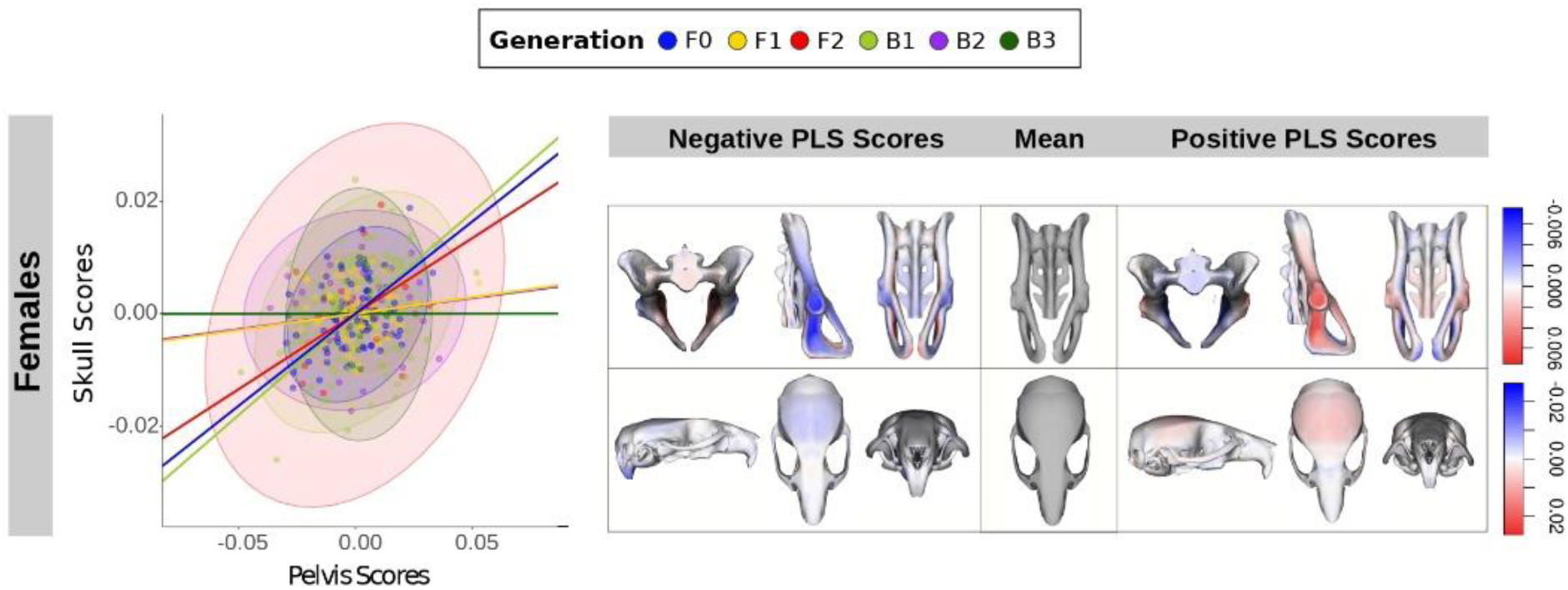
Covariance between pelvis and skull shape across generations and strains, shown for females only. Females exhibit a clear separation between generations with high versus low cephalo-pelvic covariance and a consistent association between larger, rounder neurocrania and wider, rounder pelvic canals with an open pubic symphysis. This pattern is absent in males (Supp. Fig. 7). Colored lines denote correlation coefficients derived from PLS scores of pelvis and skull blocks, calculated separately for each generation; ellipses indicate 95% confidence intervals. Shapes corresponding to positive (right) and negative (left) PLS scores are illustrated in transverse, sagittal, and coronal views. Heatmaps show expansion (red) and contraction (blue) relative to the mean shape.

### Female pelvis shape adjusts to neurocranium size only in parental generations

We expanded our investigation by analysing the relationship between female pelvis form and adult female neurocranium size as a proxy for neonatal neurocranium size (48). Procrustes regression of pelvis form conducted on non-parous F0 females was significant (*r*=0.36, *p*=0.002) and skull size accounted for 7% of pelvis shape variance. Females with large heads present, on average, a wide and round pelvic canal, wide pubic gap, short sacrum, and thin pubis. In contrast, females with a small head presented a closed pubic gap, long sacrum and slightly thicker pubic bone. However, when correcting for pelvis centroid size, the association between neurocranium size and pelvis shape was no longer significant.

Regressing pelvis form on neurocranium size in F1, F2, and backcrosses produced mixed-significance results: B1 and B2 show significant association of pelvis shape and neurocranium size (for B1 r=0.26, p=0.007; for B2 r=0.46, p=0.01), whereas F1, F2 and B3 show non-significant associations (Supp. Table 16). Pelvic morphology associated with small and large neurocranium size followed different patterns than the one observed in F0. Only in F1 the pubic region presented a difference between individuals with large and small neurocrania. F2, B1, B2 and B3 show varying degrees of sacrum length and pubic caudal elongation, but no evident differences between the opening of the pubic gap.

**Fig. 4.**
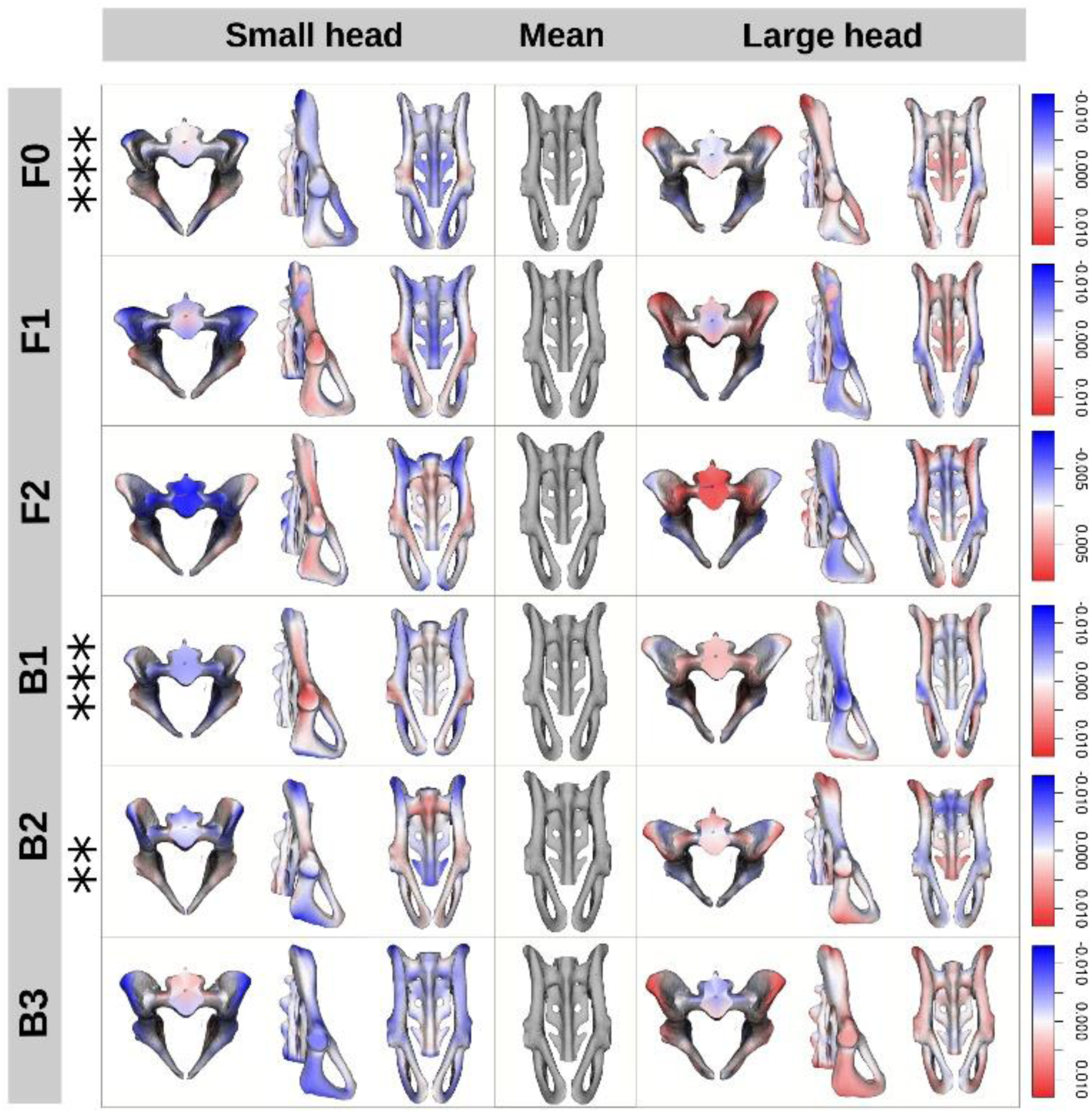
Pelvis’ heatmaps depicting the Procrustes regression of female pelvis shape on neurocranium centroid size, by generation. The association of large neurocrania and wide pelvic canal with open pubic symphysis is only present in F0 and in F1 (although non-significant in F1). This association then disappears in later generations. The morphs were obtained by warping a pelvis 3D surface onto the landmark configurations corresponding to the partial linear regression coefficients for neurocranium centroid size, controlling for body weight. Exaggerated morphs for pelvis shape are visualized in transversal, sagittal and coronal view. Heatmaps represent expansion (red) or contraction (blue) of the exaggerated morph with respect to the mean shape. Asterisks represent significance values (*: p<0.05, **: p<0.01, ***: p<0.001).

### The degree of pelvis sexual dimorphism decreases with hybridization

Sexual dimorphism in the whole sample was significant (*p*<0.001, Supp. Table 17) and accounted for 8% of the total variance in pelvis shape, while strain differences accounted for 30%, allometry for 17%, weight for 0.4% and age for 9% of shape differences. Males and females separated along both PC1 (42.4%) and PC2 (10.7%) (Supp. Fig.8). Procrustes distance between mean sex shape was the highest in F0 followed by the other generations (Fig. 5). All generations display significant sex differences, with females showing a short, dorsally tilted sacrum, thin elongated pubis, short ilia, and a spacious birth canal (Supp. Fig. 9). In F0, F1 and B1, the gap between pubic bones is more sexually dimorphic compared to F2, B2 and B3 (Fig. 5 and Supp. Fig. 9). F2 shows more sex differences in the position of the sacrum, where it is tilted dorsally compared to males. Sex differences become more subtle in the backcrosses B2 and B3. Procrustes distances were the highest between mean female and mean male shape in F0, and progressively decreased in each of the following generations (Fig. 5).

**Fig. 5.**
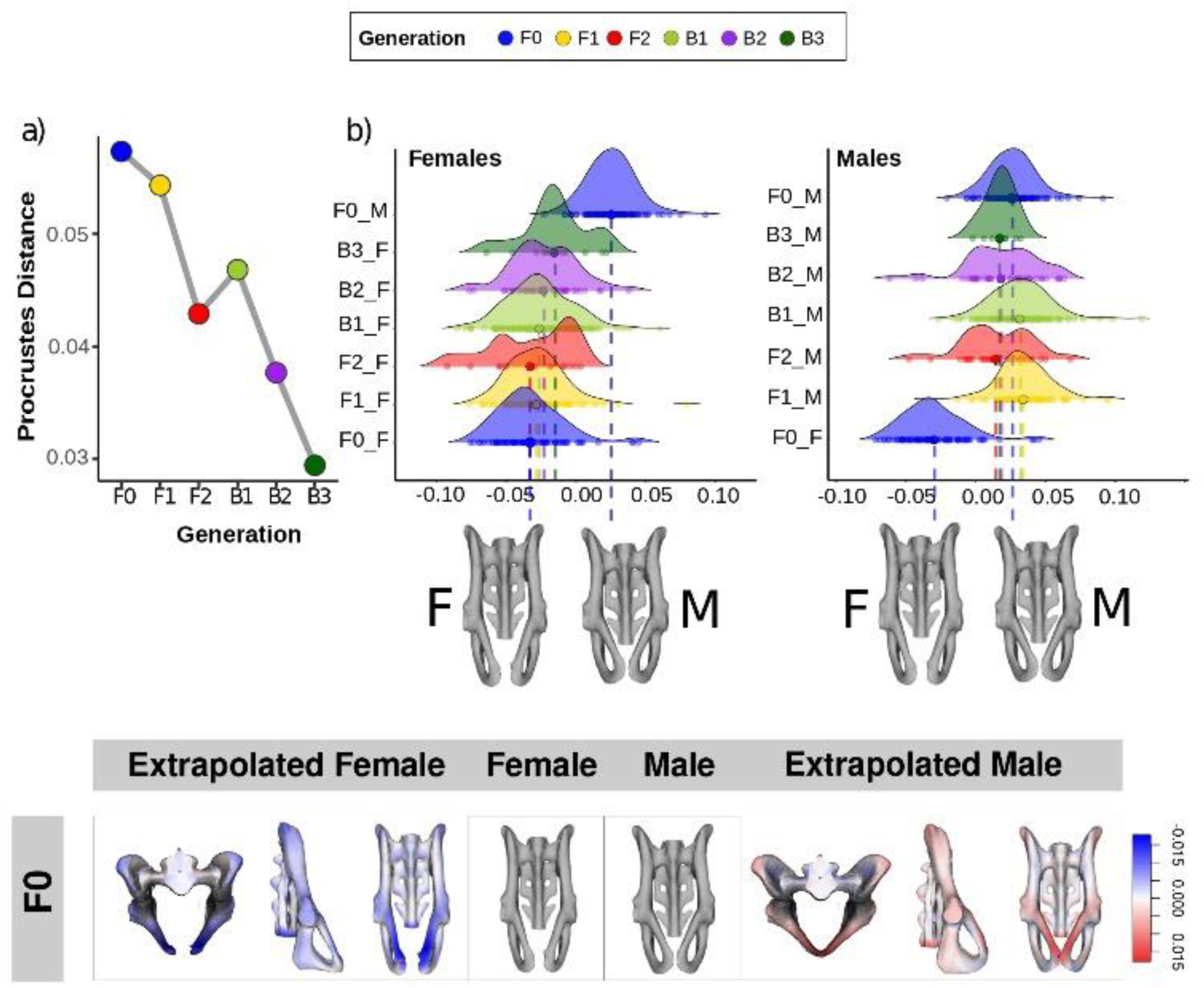
Pelvic sexual dimorphism is reduced in hybrids, primarily through partial masculinization of female pelvis shape. (*a*) Procrustes distances between male and female pelvis shapes by generation. (*b*) Ridge plot illustrating the mean female and mean male shapes in F0, along with the vector representing the shape differences between them. Pelvic shapes from individuals in the hybrids and backcross generations were subsequently projected onto this vector. Compared to the mean F0 female pelvis shape, mean female shape for hybrids and backcrosses is displaced towards the F0 male pelvis shape.

The comparison between the parental vector of sexual dimorphism and the projected values of all individuals highlighted a displacement towards the mean pelvis shape for hybrid females (Fig. 6). Hybrid females consistently display a pelvis shape closer to the male mean shape of F0, compared to F0 female pelvises. Hybrid males show mixed results, with some generation means displaying a more “feminized” pelvic shape (F2, B2, B3), while others show a “hypermasculine” shape (F1, B1) compared to the mean F0 male shape.

## Discussion

In this paper we confirmed the presence of strong skull-pelvis shape and size covariance in female mice, signatures of obstetric selection. F0 parental wild-derived populations display the highest degree of cephalo-pelvic covariation and sexual dimorphism. Reduced cephalo-pelvic covariance and reduced sexual dimorphism, and increased mismatch between neurocranial and pelvic size in hybrid generations suggest that hybridization disrupts the genetic and developmental architecture underlying obstetric integration. Therefore, hybrid females are likely to exhibit a higher risk of cephalo-pelvic disproportion and obstructed labour.

This study relies exclusively on adult morphology and therefore cannot directly determine whether cephalo-pelvic disproportion or obstructed labor occurred. Our inferences are based on morphological proxies, namely cephalo-pelvic shape and size covariation and sexual dimorphism, which are commonly used in evolutionary and biological anthropology to estimate obstetric risk, but they do not constitute direct measures of parturition outcomes (3,24,30). Consequently, the measurement of the relationship between the observed covariance patterns and actual birth difficulty remains indirect. Related to this, we used adult neurocranial size as a proxy for fetal head size (4,10,48). To assess cephalo-pelvic covariance in mice more accurately, future studies require direct measurement of morphological covariance between fetal crania and maternal pelvises.

### Hybridization leads to mismatches between cranial and pelvic size

Hybridization in wild-derived mice produced a consistent enlargement in both cranial and pelvic size in F1, a result consistent with heterosis (49–52). Later generations exhibit asymmetric size responses. Female hybrids and backcrosses show in imbalanced size response for skull and pelvis, with neurocrania growing proportionally larger than pelvises compared to F0.

This discrepancy between the pelvic size of one generation and the skull size of the next is a risk factor for feto-maternal cephalo-pelvic disproportion and obstructed labour. Because mouse neonatal head size correlates with adult head size (53), females of one generation that give birth to fetuses with disproportionately large neurocranium dimensions might encounter severe challenges during parturition. In our sample, F1, B2, and B3 neurocrania exceed the pelvic dimensions of their respective parental generations. This pattern is relevant because heterosis and in general evolutionary changes in the dimensions of the pelvis that do not follow increases in neurocranium size potentially increase the incidence of dystocia (22,40,41).

### Pelvic and cranial shape variances respond differently to hybridization

Hybridization did not affect pelvis and cranium shape equally. Pelvic morphology responds readily to hybridization, producing extreme phenotypes not observed in any of the parental strains. Hybridization frequently increases phenotypic variance and can generate extreme phenotypes, defined as trait values that fall outside the parental range (54–56). The emergence of such phenotypes can inflate overall shape variance and expand the morphospace available to selection in hybrid generations. Skull morphology didn’t respond with the same increase in variance. Hybrids and backcrosses presented reduced variance in skull shape and a mostly intermediate morphology to that of the parents. However, when controlling for between-strain mean shape differences, new hybrid phenotypes emerged. This indicates that once divergent mean shapes are established between strains during species radiation, the within-strain skull covariance matrix is largely similar. This asymmetry between traits’ response to hybridization indicates that different traits might have different degrees of genetic architecture stability. In our case, pelvises might possess a more unstable and versatile genetic architecture compared to craniofacial genetic architecture. Reduced variance in hybrids skull morphology can limit the morphospace available to selection and consequently influence hybrids’ integration between functionally related traits. Moreover, reduction of functional integration in favor of developmental modularity has been recorded for other mammals (57). In this case, pelvis shape variance, and not functional obstetric integration, represents the main driving force for either the consolidation of cephalo-pelvic covariation, or its reduction, depending on how the variance-covariance matrix of pelvis and skull align.

### Cephalo-pelvic integration is female-specific

Functionally codependent traits must covary in a coordinated manner to maintain proper function (9). The appropriate covariance between adult skull and pelvic form translates to covariance between fetus’ head and mother’s pelvic form, minimizing the damage caused by cephalo-pelvic disproportion and obstructed labour. The presence of significant covariation between skull and pelvic shape in adult mice suggests that cephalo-pelvic covariation might be repeatedly selected in mammals to alleviate dystocia. In females derived from wild mouse populations, we found that wider rounder birth canals are associated with larger and rounder neurocrania, a pattern shared with humans and other primates (23–25,31). This intriguing pattern of female-specific morphological covariation might involve sex-specific processes that regulate maternal pelvis development, adult head development and possibly fetal head development. The dependence of this pattern on size however suggests that the morphological accommodation of neurocranial size is in part mediated by overall body size. The association between rounder neurocrania and rounder birth canals was not detected in males, likely justified by the absence of obstetric selection (28,58).

### Pelvis and cranium covariance are reduced with hybridization

Contrary to F0 parental populations, we observed that cephalo-pelvic covariance was variously reduced in some hybrid generations. It has been proposed that functional integration of obstetrically relevant traits in females is obtained through developmental and genetic integration (4). Covariance between traits requires variance in pleiotropic developmental processes and alleles (59). Hybridization often causes shifts in trait covariance due to genetic introgression, establishment of new dominance patterns, and transgressive segregation (60,61). The results are emergence of novel pelvis or cranial shapes, extreme phenotypes (larger neurocrania, smaller pelvises), or reassortment of phenotypic traits (mismatch between pelvis and cranial shape) (35–38).

Studies of between-trait shape covariance show that hybridization often weakens functional integration in F1 and F2. Consistent with this, our F1 hybrids exhibit intermediate shapes but disproportionately enlarged skulls relative to both parents and pelvic size, reducing skull-pelvis integration. F2 individuals display high pelvic shape variance, which inflates correlation coefficients with skull shape, but also high phenotypic dispersion. This pattern aligns with observed changes in trait variance-covariance matrices for F2 populations (62). Accordingly, we observe limited morphological functional integration between pelvic morphology and neurocranial size. Because selection has not yet removed mismatched phenotypes, scatter around the main axis of variation remains high, and covariance structure is likely unstable relative to parental populations (63). Backcrosses B2 and B3, despite retaining a high proportion of parental genomic content (and thus expected to regress toward parental levels of cephalo-pelvic covariance) show some of the lowest covariance values. The mechanisms behind the loss of covariance in backcrosses warrant further investigation.

### Hybridization reduces pelvic sexual dimorphism

Hybridization also affects sexual dimorphism: while we observed a high degree of sexual dimorphism in parental F0 generations, subsequent generations presented decreased sex differences. The mechanisms that lead to reduced or increased sexual dimorphism in hybrids are similar to the processes that reduce or increase trait variation and covariation. Sexual dimorphism emerges when the two sexes are driven by selection to different sex-specific optimal phenotypes (64). Complex morphological traits like sexual dimorphism are highly polygenic, and their genetic architecture evolves independently in different populations over time. Different populations might achieve the final optimal female and male pelvic form based on different additive and non-additive genetic variance. When two parental populations with differing underlying genetic variation and different sex-specific developmental pathways interbreed, the resulting hybrids might show dampening (or intensifying) sexual dimorphism (45). Interestingly, in our study the reduction of the degree of pelvic sexual dimorphism is constant and does not depend on the level of variance of or covariance with skull shape or neurocranium size. While some male hybrids showed both a “feminization” (F2, B2 and B3), or “hyper-masculinization” (F1, B1) of their pelvic shape, hybrid females only tended to show a gradual “masculinization” of their pelvis shape along the sexual dimorphism vector. This displacement towards the mean shape shown by female hybrids is not explained by parity status nor by mean age differences between the generations, consistently with other studies (65). The reduction of sexual dimorphism for obstetrically selected traits and a “masculinized” pelvic phenotype consisting of a narrower birth canal, could negatively impact childbirth.

## Conclusion

In this paper we proved that CPD and cephalo-pelvic covariance are not unique to humans or primates, but are shared with several other mammals including the common house mouse. The presence of such covariance in a quadrupedal mammal with small feto-maternal size ratio suggests that obstetric selection acting on covariation between head and pelvic morphology is not exclusively tied to the evolution of encephalization, nor to the transition to bipedality in *Homo*. We further showed that hybridization modifies the variance-covariance structure of head and pelvic shape and size, and alters both the magnitude and the pattern of sexual dimorphism. These patterns are consistent with differences among the parental strains in the genetic architecture underlying cephalo-pelvic covariation and sexual dimorphism. When these architectures are combined through hybridization, the associations that structure cephalo-pelvic covariation and sex-specific morphology become diffused. The mechanistic bases of these changes remain to be investigated.

These results are important because hybridization is common in evolution, including the evolution of the human lineage. Hybridization between *Homo sapiens*, Denisovans and Neanderthals happened in Eurasia several times, with potential effects on the morphology of skull and pelvis of modern humans (66–70). Despite the small number of available hominid fossils, studies have speculated that Neandertals used to give birth to large-headed, heavy neonates, and that despite their wide birth canal, this process was just as painful and complicated as it is in modern humans (71). Modern humans and Neanderthals cranial and pelvic form has been subjected to the action of climate adaptation and neutral divergent selection, creating diverse morphologies and patterns of sexual dimorphism (71–74). Hybridization between two species of *Homo* with difficult childbirths and cephalo-pelvic proportions could have increased temporarily the incidence of cephalo-pelvic disproportion, causing intense transient obstetric selection in these human hybrid populations (75,76).

Finally, reduced cephalo-pelvic covariance in hybrid mice could in part explain trends in modern-day cephalo-pelvic disproportion incidence. Human populations display a variety of geographically structured birth canal and cranial shapes, with some specific traits like neurocranial shape reflecting neutral genetic distances (72,77). Cephalo-pelvic covariation might be structured in a similar way, with different populations evolving patterns of covariation that follows specific anatomical adaptations. Future studies might help address whether population differences in human cephalo-pelvic morphological covariation follow a similar pattern to what we observed for mouse hybrid populations.

## Acknowledgements

This work was supported by CIHR Foundation Grant 159920, a Canada First Research Excellence Program Grant (One Child Every Child, CFREF 2022-00015), NSERC Discovery Grant 238992-23 and the Canadian Foundation for Innovation, Innovation grant #36262 to BH. This project was also funded by the National Research Foundation of South Africa (Grant no. 81824) and the National Research Foundation of South Africa Grant no. AOP240509218040 to RRA. EZ is supported by the ACHRI Graduate Fellowship of the University of Calgary, and the Konrad Lorenz Institute Writing Fellowship (Konrad Lorenz Institute, Vienna, Austria).

## Methods

### Mouse Model

All mice were housed and bred at the University of XXX in accordance with approved animal care protocols from both the University of XXX and the University of XXX (AC1-0210 and 2012V56RA, respectively). Mice were derived from four parental strains of wild mice: *Mus musculus spretus* (SPRET), *M. m. castaneus* (CAST), *M. m. musculus* (CZE) and *M. m.domesticus* (WSB). The wild parental strains were captured away from known interbreeding zones. The four initial strains of parental mice were crossed to produce five different interspecific F1 hybrids. Two F2 hybrid strains resulted from inbreeding of F1 hybrids. Male infertility between males of the hybrid WSBXCZE prevented breeding of WSBXCZE_F2. Backcrosses were also bred for six B1 backcrosses result of the cross between an F1 and a parental strain, two B2 backcrosses resulted from a cross between a F2 and a parental strain, and two B3 crosses between a B1 backcross and a F0 (Table1, Supp. Materials). Hybrids derived from WSBXCZE and hybrids derived from SPRET were sterile, therefore no F2, B1, B2 or B3 hybrids could be produced from these strains. The final sample after removing outliers comprised 802 mice from 6 generations and 21 strains (Supp. Fig 6 and Supp. Table 1).

### Pelvis and Skull morphology

For each mouse, we collected 3D skull and pelvis morphology using x-ray micro-computed tomography (µCT). Scanning was done at the University of XXX with either a Scanco vivaCT 40 µCT scanner or a Scanco vivaCT 80 µCT scanner (Scanco Medical, Brüttisellen, Switzerland) at 0.035 mm voxel dimensions at 55kV and 72-145 μA. Given the high number of specimens, manual landmarking was replaced by an atlas-based automated landmarking pipeline developed at the Hallgrímsson lab, and explained in detail in Devine et al 2020 and in the corresponding MusMorph Github repository (https://github.com/jaydevine/MusMorph/tree/main/Preprocessing) (78).

After initial reorientation of skulls and individual pelvic bones, we created one reference atlas for skull and one for each pelvic bone by spatially normalizing the microCT images with a group-wise registration workflow using the MNI Display software (Montreal Neurological Institute, http://bic-mni.github.io/man-pages/). The atlas generation scripts are available in the MusMorph Github repository (https://github.com/jaydevine/MusMorph/tree/main/Processing, HiRes_Atlas.py or LoRes_Atlas.py). We then manually segmented the skull average to specify the bony structures from the surrounding soft tissue, using the Segmentation toolkit in MINC (http://bic-mni.github.io/man-pages/). After rendering the average atlas in MINC using the built-in display tools, we manually placed a set of anatomical landmarks on skull surface, using the MNI Display software (336 landmarks, real landmarks = 94, curve semilandmarks=242) (Fig 2 Supp. Materials).

Each image was registered to their atlas and the landmarks backpropagated using the Bash and Python script HiRes_Pairwise.py and Label_Propagation.py at (https://github.com/jaydevine/MusMorph/tree/main/Processing).

High variation in pelvis morphology prompted a slightly different phenotyping strategy. The three main bones of the female pelvis (right os coxae, left innominate and sacrum) often display great variation in position due to loose articulations. Hence, we decided to perform the automated landmarking algorithm on the three bones separately, and subsequently reconstruct all pelvises virtually. The “noise” due to random rotation of the os coxae on the flexible sacroiliac joint was reduced by reflecting paired bilateral landmarks across a least-square-fitted midsagittal plane define by the central sacral landmarks in R (R version 4.5.0, R core team). For each pelvis, we obtained two landmark configurations (an original reconstructed one and its mirrored counterpart) that were then superimposed and averaged. The final pelvis landmark scheme included 213 landmarks (real landmarks =30, curve semilandmarks=183, Fig 1 Supp. Materials).

### Statistical analyses

#### Shape data acquisition

We standardized pelvis, skull and neurocranium landmark data for position, rotation and size using Generalized Procrustes Analysis or GPA (79) in the package *geomorph* in R (Adams et al., 2013, R Core Team) We plotted outliers using the function plotOutliers in *geomorph* and manually inspected specimens that exceeded the upper quartile threshold. We conducted this operation separately for pelvises and skulls and removed specimens that were damaged or either outliers for skull or pelvis.

#### Pelvis size and skull size comparison across generations

To test for possible size mismatch between generations, we measured mean pelvis size and neurocranium size (instead of whole skull size) for every generation using their centroid size. The neurocranium was chosen instead of the whole skull because it grows linearly throughout the life of a mouse and presents high heritability (53), providing a more accurate proxy for fetal head size.

We computed centroid size for pelvis and skull separately using the function cSize from the R *Morpho* package (81) from the raw, unaligned coordinates. Strain-level centroid size was obtained as the mean centroid size of individuals belonging to that strain. These strain means were then expressed as percent differences relative to the F0 reference, where the F0 baseline was defined as the average centroid size of all F0 individuals. This was repeated for pelvis and skull centroid sizes of each strain. Centroid size differences between generations were computed via ANOVA. Pairwise significances were calculated with the function pairwise.t.test (R package *stats,* R version 4.5.0, R core team) for Welch two-sample, two-sided pairwise t-tests between all the generations, applying Benjamini-Hochberg correction for multiple comparisons.

#### Pelvis shape and skull shape variance

After obtaining the landmark configurations for pelvis and skull from our automated landmarking pipeline, we performed a Generalized Procrustes Analysis (GPA) to remove the effect of translation, rotation and scaling. The resulting Procrustes-aligned landmarking configurations were then used for each subsequent analysis. Pelvis and skull shape variance was visualized by computing a Principal Component Analysis using the function gm.prcomp from the package *geomorph.* PCA was first calculated on generation-mean-corrected pelvis and skull landmark configurations, and then on strain-mean-corrected configurations.

#### Pelvis and skull shape covariance

We conducted two-block partial least squares (2B-PLS) analyses between Procrustes-aligned pelvic and skull shape data using *geomorph*, controlling for sex, parity (females), and strain. Separate analyses were performed for females and males. For each generation, correlations between pelvis and skull PLS scores were computed with cor.test in *stats*. We visualized pelvis and skull shapes associated with positive and negative PLS axes by multiplying the vector of shape change by a multiple of the standard deviation of the vector length and adding or subtracting it to the consensus shape of the sample. We then generated both morphs and heatmaps to visualize these shapes and their differences, using the function meshDist from the package *Morpho* (Schlager S et al, 2017). For each 2B-PLS analysis block, we extracted the skull and pelvis scores and grouped them by generation. We then calculated the correlation between skull and pelvis scores for each generation using the function cor.test from the package *stats (*R version 4.5.0, R core team). To calculate the dispersion around the correlation axis, we calculated the Mean Square Error by squaring the residuals of the regression calculated for skull and pelvis scores, separately for each generation.

#### Pelvis shape and neurocranium size covariance

Neurocranium coordinates were extracted by subsetting 150 landmarks from the skull coordinates. For a reference see Supp. Fig. 3. We calculated neurocranium centroid size using the function cSize from package *Morpho* on the raw un-aligned neurocranium coordinates. We then conducted Procrustes shape regression of pelvis form (shape + size) on the standardized neurocranium size in F0 strains (CAST, WSB, CZE and SPRET, 89 females) pooled together, correcting for strain and body weight for both. We visualized the pelvic morphology associated with relatively small and large neurocranial sizes by adding ten times the vector of pelvis shape change, scaled to its standard deviation, to the mean configuration. To calculate the correlation between pelvis shape scores and neurocranium residuals after correction for body weight and strain, we used the function *cor.test* in R.

#### Sexual dimorphism

Pelvic sexual dimorphism was tested using Procrustes regression of pelvis shape on sex across 803 Procrustes-aligned specimens spanning parental, hybrid, and backcross generations; one B3 strain with a single female was excluded. Shape was regressed on sex controlling for allometry using body weight, using procD.lm in *geomorph*. The magnitude of sexual dimorphism was quantified as the Procrustes distance between mean male and female shapes per generation, and significance assessed by permutations within-generation. Residual shapes were summarized by PCA for visualization, and exaggerated morphs are produced by multiplying by two the vector of male-female shape and adding or subtracting it from the mean female and male shape. Generation-specific dimorphism was further projected onto the F0 sex-difference vector to compare magnitudes relative to the parental baseline.

