## Supplementary_figures_tables for "Cephalo-pelvic covariation and sexual dimorphism are disrupted in hybrid mice: implications for the human obstetrical dilemma"

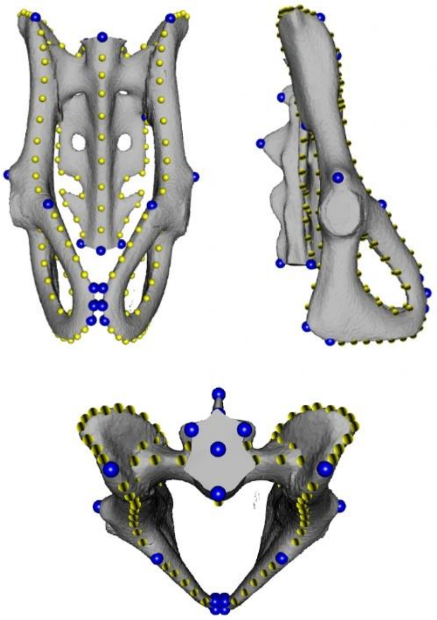

Supp. fig. 1. Pelvis landmark scheme. Blue = fixed landmarks. Yellow = semilandmarks.

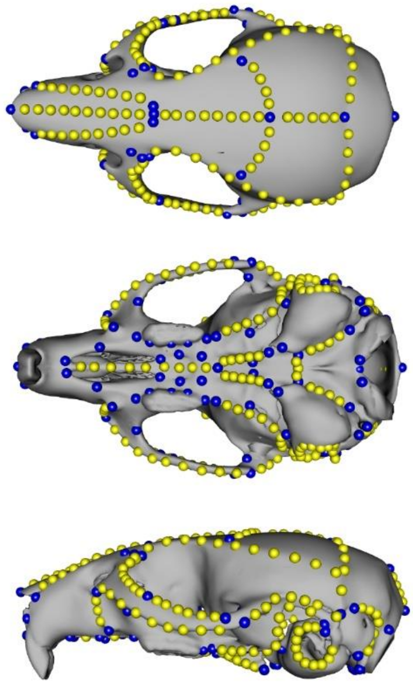

Supp. fig. 2. Skull landmark scheme. Blue = fixed landmarks. Yellow = semilandmarks.

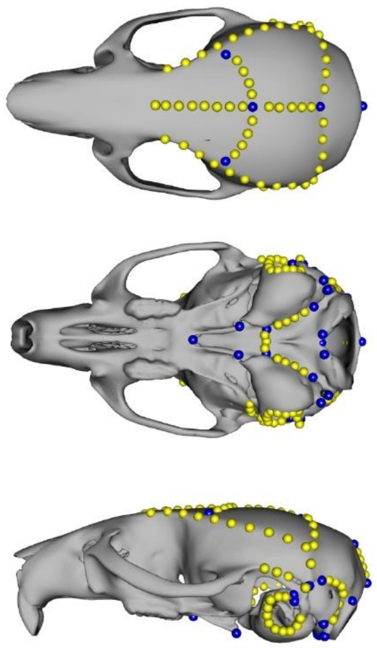

Supp. fig. 3. Neurocranium landmark scheme. Blue = fixed landmarks. Yellow = semilandmarks.

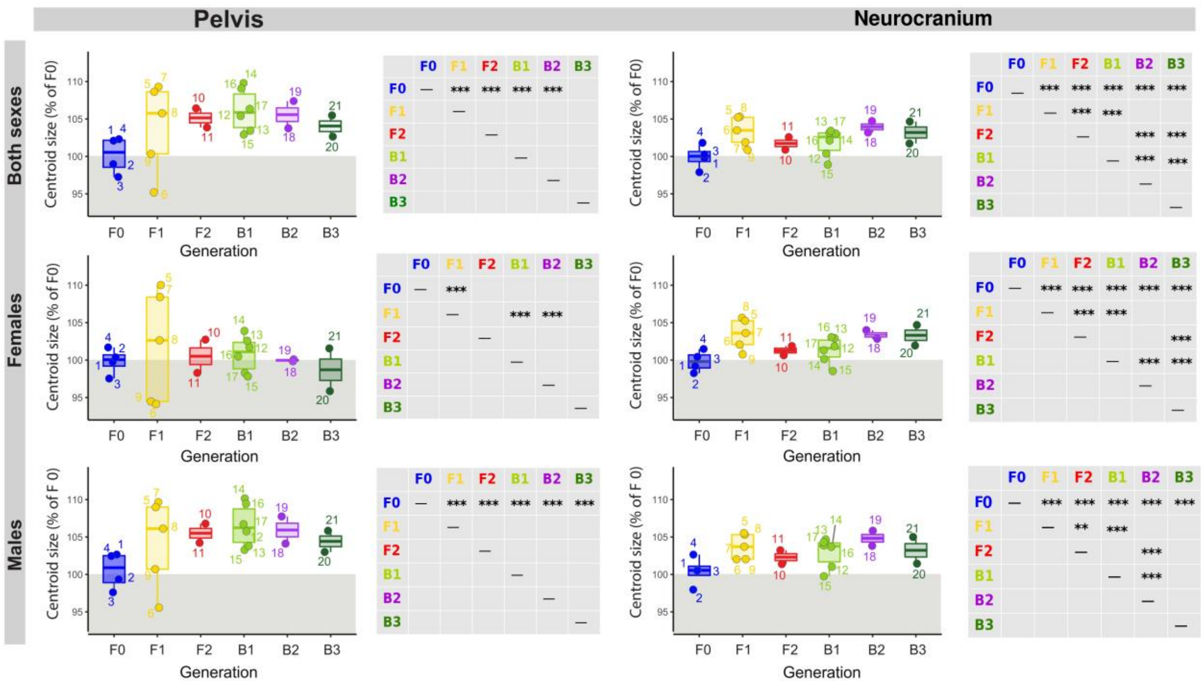


Supp. fig. 4. Mean pelvis and neurocranium centroid size for each generation. The values are expressed as percentage of the mean centroid size of F0. Each dot represents the mean centroid size for each strain. Significant differences between generations are reported in the grey grids as asterisks (p<0.05*, p<0.01**, p<0.001***).

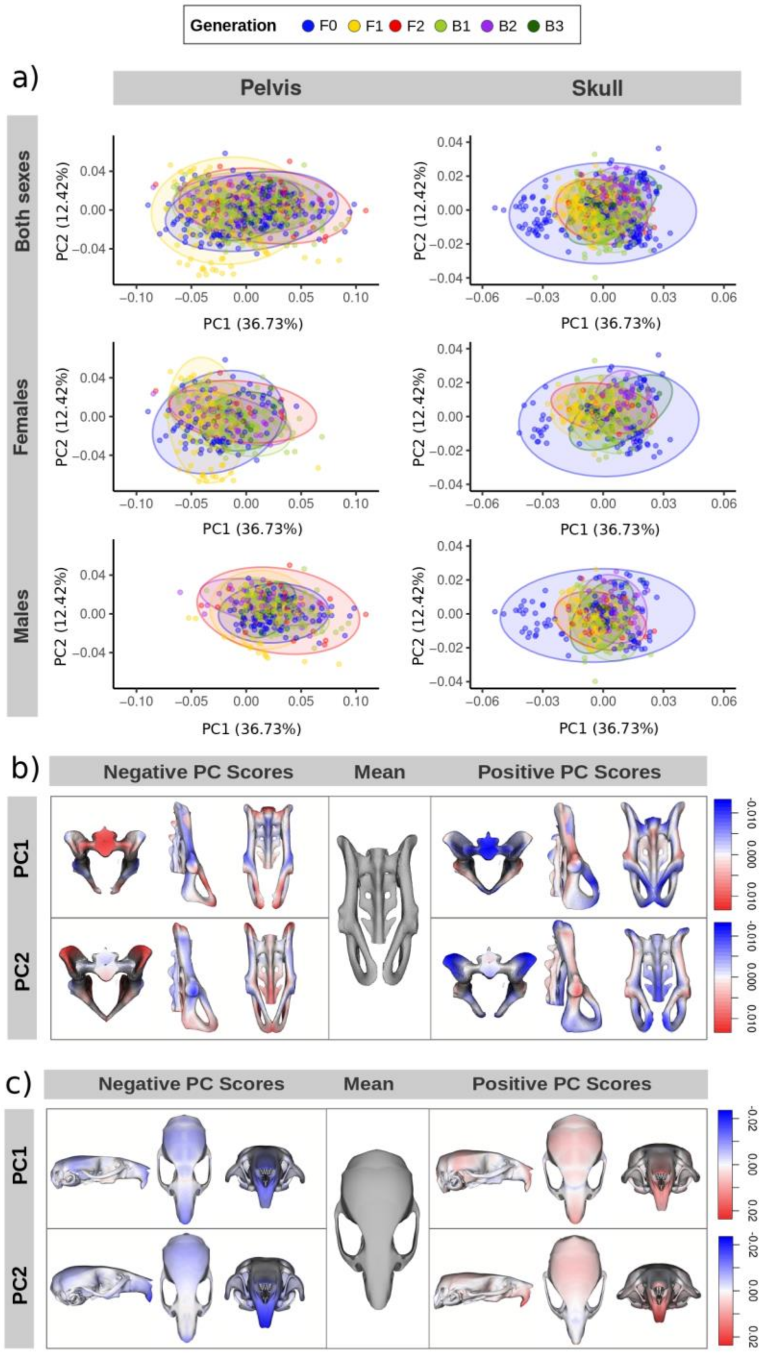


Supp. fig. 5. (a) PCA scatter plots for pelvis and skull for both sexes, only females and only males, mean centered on Generation. (b) Morphs for pelvis and (c) skull shape along PC1 and PC2. Morphs were generated by scaling PC loading vectors by 4× their respective eigenvalues (pelvis) or by 5× (PC1) and 10× (PC2) eigenvalues (skull). Heatmaps indicate expansion (red) and contraction (blue) relative to the mean.

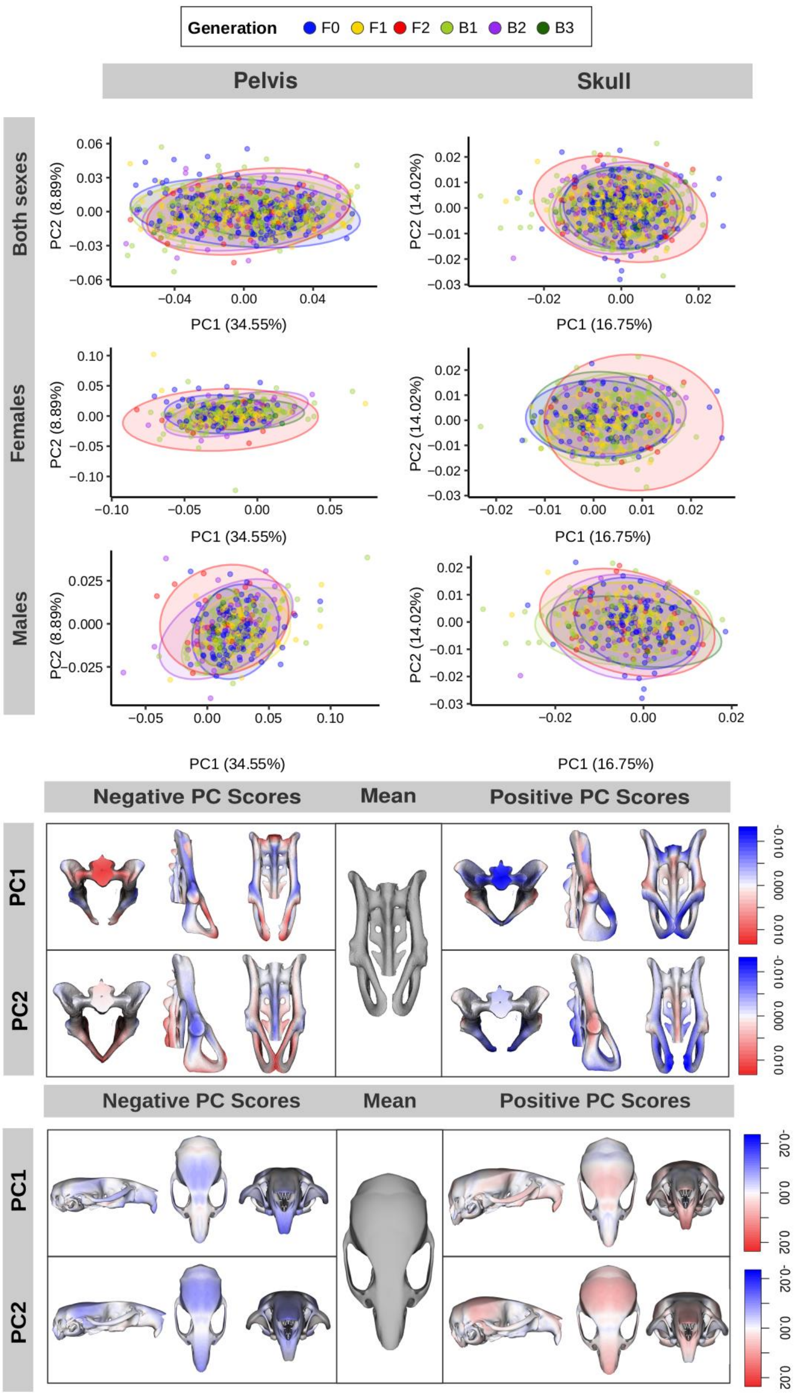


Supp. fig. 6. (a) PCA scatter plots for pelvis and skull for both sexes, only females and only males, mean centered on strain. (b) Morphs for pelvis and (c) skull shape along PC1 and PC2. Morphs were generated by scaling PC loading vectors by 4× their respective eigenvalues (pelvis) or by 5× (PC1) and 10× (PC2) eigenvalues (skull). Heatmaps indicate expansion (red) and contraction (blue) relative to the mean.

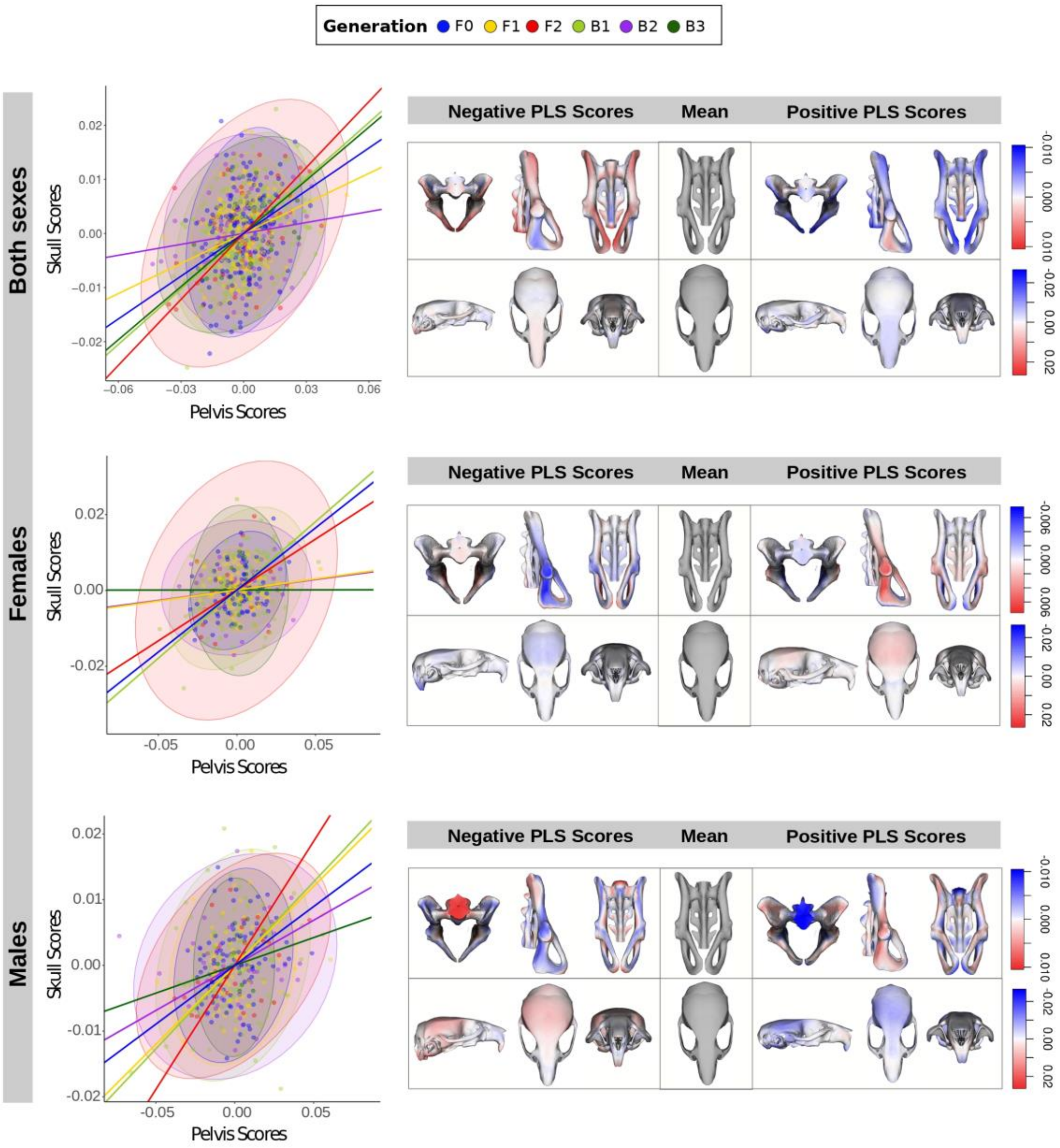


Supp. fig. 7. Covariance between pelvis and skull shape for both sexes (top), females only (middle) and males only (bottom). Females exhibit a clear separation between generations with high versus low cephalo-pelvic covariance and a consistent association between larger, rounder neurocrania and wider, rounder pelvic canals with an open pubic symphysis. This pattern is absent in males (Supp. Fig. 7). Colored lines denote correlation coefficients derived from PLS scores of pelvis and skull blocks, calculated separately for each generation; ellipses indicate 95% confidence intervals. Shapes corresponding to positive (right) and negative (left) PLS scores are illustrated in transverse, sagittal, and coronal views. Heatmaps show expansion (red) and contraction (blue) relative to the mean shape.

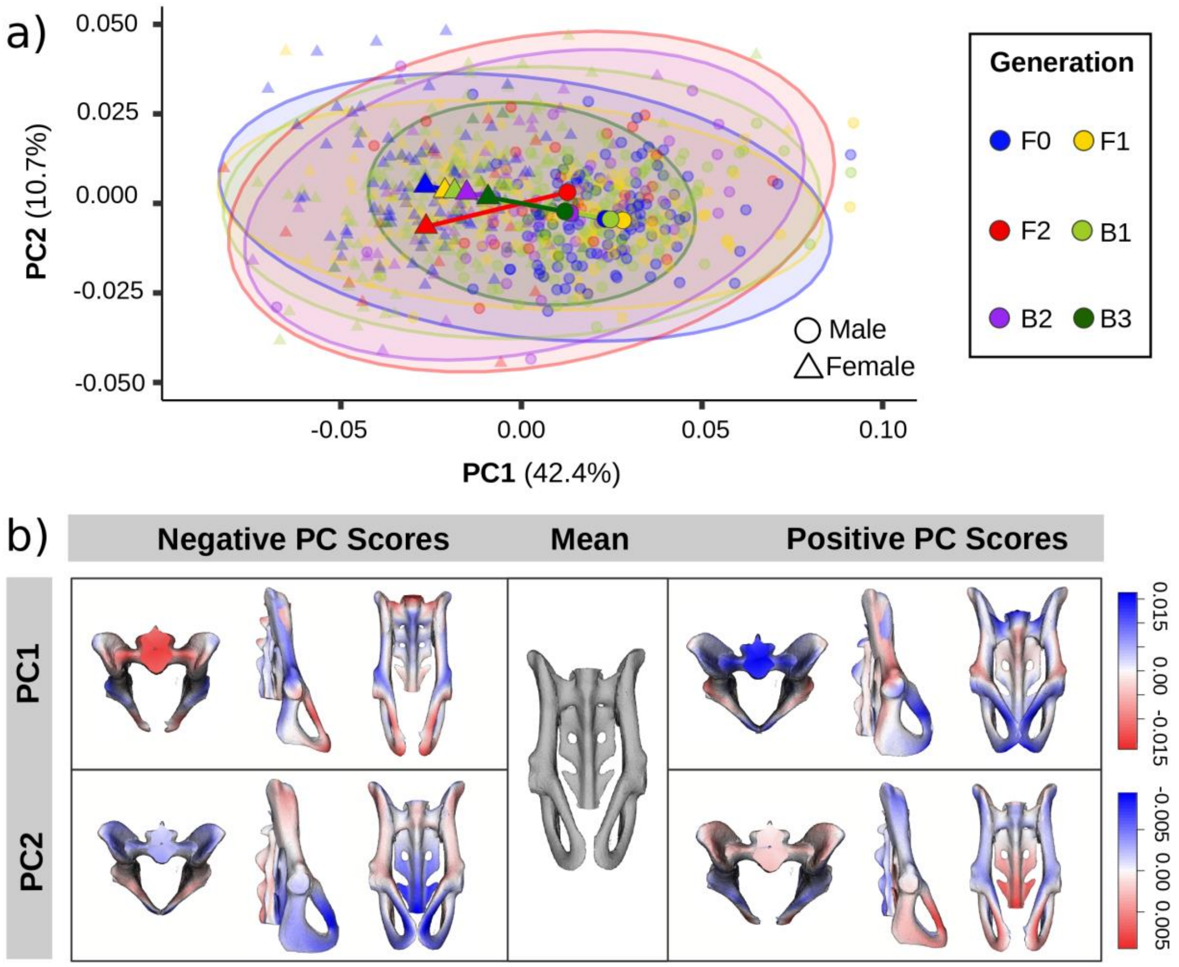


Supp. fig. 8. (a) PCA of pelvis shape corrected for strain, age, and size, with vectors indicating mean sex differences along PC1 and PC2 for each generation; colored ellipses denote 95% confidence intervals. (b) Mean pelvis shape (grey) with exaggerated morphs along PC1 and PC2 in superior, lateral, and anterior views; morphs were generated by scaling loading vectors by 4× their respective eigenvalues. Heatmaps indicate expansion (red) and contraction (blue) relative to the mean.

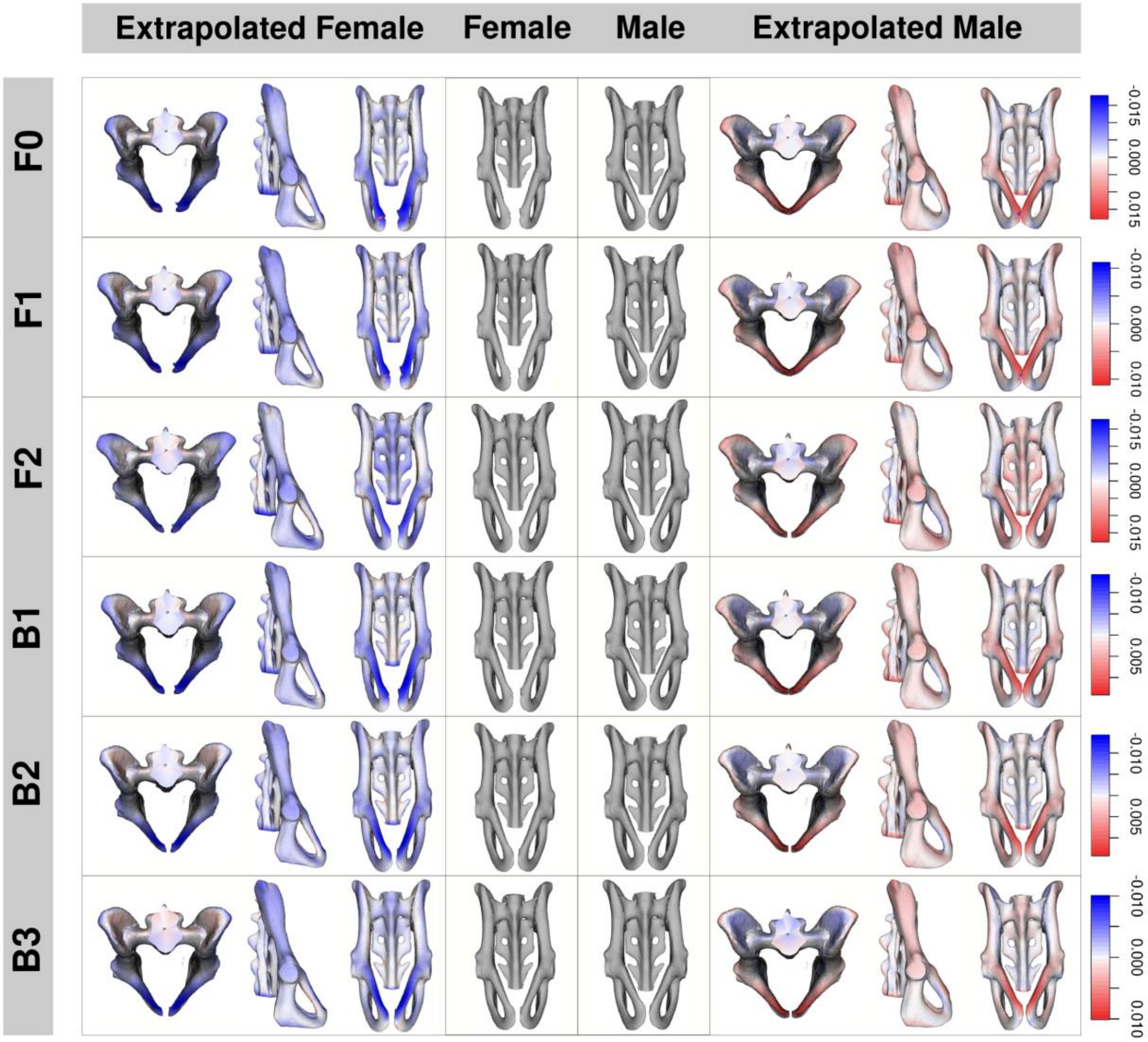


Supp. fig. 9. Sex differences in pelvic shape by generation. Mean female and mean male shape (grey) and exaggerated morphs of sex differences for female pelvis shape (left) and male pelvis shape (right). These morphs were calculated by multiplying by a factor of two the vector of shape differences between mean male from mean female pelvis shape. We then added this vector to the mean female pelvis shape to obtain the exaggerated female pelvis shape, and subtracted it from the mean male pelvis shape to obtain the exaggerated male pelvis shape. Heatmaps represent expansion (red) or contraction (blue) of the exaggerated morph with respect to the mean shape.

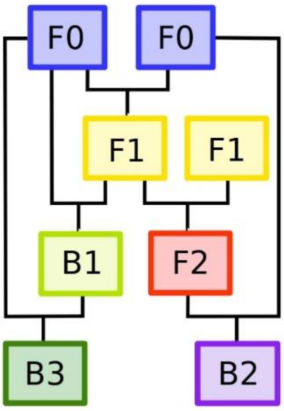


Supp. fig. 10. Mouse breeding scheme.

| Code | Strain | Generation | Parent 1 | Parent 2 | N | Females | Males |
|---|---|---|---|---|---|---|---|
| 1 | CAST | F0 | CAST | CAST | 49 | 21 | 28 |
| 2 | CZE | F0 | CZE | CZE | 43 | 18 | 25 |
| 3 | SPRET | F0 | SPRET | SPRET | 50 | 20 | 30 |
| 4 | WSB | F0 | WSB | WSB | 49 | 29 | 20 |
| 5 | WSBXCZE | F1 | CZE | WSB | 48 | 29 | 19 |
| 6 | WSBXSPRET | F1 | WSB | SPRET | 35 | 19 | 16 |
| 7 | CASTXCZE | F1 | CAST | CZE | 46 | 25 | 21 |
| 8 | CASTXWSB | F1 | CAST | WSB | 48 | 25 | 23 |
| 9 | CASTXSPRET | F1 | CAST | SPRET | 9 | 8 | 1 |
| 10 | CASTXCZE_F2 | F2 | CASTXCZE | CASTXCZE | 20 | 7 | 13 |
| 11 | CASTXWSB_F2 | F2 | CASTXWSB | CASTXWSB | 20 | 6 | 14 |
| 12 | WSBXCZE_CZE | B1 | WSBXCZE | CZE | 49 | 26 | 23 |
| 13 | WSBXCZE_WSB | B1 | WSBXCZE | WSB | 50 | 30 | 20 |
| 14 | CASTXCZE_CAST | B1 | CASTXCZE | CAST | 47 | 28 | 19 |
| 15 | CASTXCZE_CZE | B1 | CASTXCZE | CZE | 37 | 21 | 16 |
| 16 | CASTXWSB_CAST | B1 | CASTXWSB | CAST | 48 | 30 | 18 |
| 17 | CASTXWSB_WSB | B1 | CASTXWSB | WSB | 49 | 25 | 24 |
| 18 | CAXW_CAXW_W | B2 | CASTXWSB_F2 | WSB | 44 | 19 | 25 |
| 19 | CAXW_CAXW_CA | B2 | CASTXWSB_F2 | CAST | 39 | 20 | 19 |
| 20 | CA _CAXW_CA | B3 | CASTXWSB_CA | CAST | 4 | 1 | 3 |
| 21 | W _CAXW_CA | B3 | CASTXWSB_CA | WSB | 19 | 10 | 9 |

**Table 1** summary of mouse strains included in the sample, including generation, parental strain derivation, and sample size.

| Generation | Both sexes | Females | Males |
|---|---|---|---|
| F0 | 87.4 | 88.6 | 86.3 |
| F1 | 91.5 | 92.0 | 90.8 |
| F2 | 90.2 | 89.2 | 90.7 |
| B1 | 90.4 | 89.5 | 91.6 |
| B2 | 89.8 | 88.6 | 90.9 |
| B3 | 89.7 | 88.8 | 90.8 |

**Table 2** Pelvis centroid size by generation.

| Generation | Both sexes | Females | Males |
|---|---|---|---|
| F0 | 62.5 | 62.5 | 86.3 |
| F1 | 65.0 | 65.0 | 90.8 |
| F2 | 63.6 | 63.2 | 90.7 |

| | | | |
|---|---|---|---|
| B1 | 63.7 | 63.4 | 91.6 |
| B2 | 64.9 | 64.6 | 90.9 |
| B3 | 65.1 | 65.0 | 90.8 |

**Table 3** Neurocranium centroid size by generation.

| Strain | Both sexes | Females | Males |
|---|---|---|---|
| CA_CAXW_CA | 85.84 | 84.94 | 88.57 |
| CAST | 88.32 | 88.39 | 88.27 |
| CASTXCZE | 95.27 | 96.07 | 94.31 |
| CASTXCZE_CAST | 93.15 | 92.07 | 94.74 |
| CASTXCZE_CZE | 87.83 | 87.09 | 88.81 |
| CASTXCZE_F2 | 91.55 | 91.04 | 91.83 |
| CASTXSPRET | 84.03 | 83.71 | 86.58 |
| CASTXWSB | 91.10 | 90.95 | 91.27 |
| CASTXWSB_CAST | 91.63 | 90.13 | 94.13 |
| CASTXWSB_F2 | 88.87 | 87.10 | 89.63 |
| CASTXWSB_WSB | 89.16 | 86.67 | 91.76 |
| CAXW _CAXW_CA | 90.63 | 88.70 | 92.67 |
| CAXW_CAXW_W | 89.07 | 88.47 | 89.53 |
| CZECHI | 86.97 | 88.93 | 85.41 |
| SPRET | 84.92 | 86.43 | 83.92 |
| W_CAXW_CA | 90.48 | 90.00 | 91.02 |
| WSB | 89.27 | 90.11 | 88.10 |
| WSBXCZE | 96.01 | 97.49 | 93.74 |
| WSBXCZE_CZE | 89.14 | 89.04 | 89.26 |
| WSBXCZE_WSB | 90.92 | 90.90 | 90.95 |
| WSBXSPRET | 82.83 | 83.41 | 82.15 |

**Table 4** Pelvis centroid size by strain.

| Strain | Both sexes | Females | Males |
|---|---|---|---|
| CA_CAXW_CA | 63.56 | 63.69 | 63.16 |
| CAST | 62.34 | 61.97 | 62.63 |
| CASTXCZE | 64.66 | 64.73 | 64.58 |
| CASTXCZE_CAST | 63.81 | 63.30 | 64.56 |
| CASTXCZE_CZE | 61.80 | 61.56 | 62.11 |
| CASTXCZE_F2 | 63.04 | 62.88 | 63.13 |
| CASTXSPRET | 63.02 | 62.96 | 63.53 |
| CASTXWSB | 65.80 | 66.01 | 65.58 |
| CASTXWSB_CAST | 64.45 | 64.31 | 64.68 |

| | | | |
|---|---|---|---|
| CASTXWSB_F2 | 64.09 | 63.65 | 64.28 |
| CASTXWSB_WSB | 64.40 | 63.64 | 65.19 |
| CAXW _CAXW_CA | 65.43 | 64.98 | 65.91 |
| CAXW_CAXW_W | 64.48 | 64.26 | 64.65 |
| CZECHI | 61.17 | 61.39 | 60.99 |
| SPRET | 62.65 | 62.75 | 62.58 |
| W_CAXW_CA | 65.40 | 65.40 | 65.39 |
| WSB | 63.62 | 63.40 | 63.91 |
| WSBXCZE | 65.74 | 65.76 | 65.71 |
| WSBXCZE_CZE | 62.72 | 62.56 | 62.90 |
| WSBXCZE_WSB | 64.62 | 64.40 | 64.96 |
| WSBXSPRET | 63.66 | 63.78 | 63.52 |

**Table 5** Neurocranium centroid size by strain.

| **Effect** | **Df** | **$R^2$** | **F** | **Z** | **p** |
|---|---|---|---|---|---|
| Sex | 1 | 0.155 | 241.51 | 5.60 | **<0.001** |
| Strain | 20 | 0.304 | 23.74 | 14.93 | **<0.001** |
| Parity | 1 | 0.009 | 14.70 | 4.65 | **<0.001** |
| Residuals | 779 | 0.499 | – | – | – |

**Table 6** Procrustes MANOVA testing the independent effects of sex, strain, and parity on Procrustes aligned pelvis coordinates. Significance assessed by residual randomization with 1,000 permutations. Type III sums of squares. Pelvic shape showed a strong effect of sex ($R^2$ = 0.155), substantial inter-strain differences ($R^2$ = 0.304), and a small but significant parity effect ($R^2$ = 0.009), all p =0.001.

| **Effect** | **Df** | **$R^2$** | **F** | **Z** | **p** |
|---|---|---|---|---|---|
| Sex | 1 | 0.021 | 31.66 | 7.85 | **<0.001** |
| Strain | 20 | 0.447 | 33.26 | 14.19 | **<0.001** |
| Parity | 1 | 0.005 | 7.89 | 4.56 | **<0.001** |
| Residuals | 779 | 0.524 | – | – | – |

**Table 7** Procrustes MANOVA testing the independent effects of sex, strain, and parity on neurocranial shape. Significance assessed by residual randomization with 1,000 permutations. Type III sums of squares. Neurocranial shape variation was dominated by strain ($R^2$ = 0.447), with smaller but significant effects of sex ($R^2$ = 0.021) and parity ($R^2$ = 0.005), all p = 0.001.

| Effect | Df | $R^2$ | F | Z | p |
|---|---|---|---|---|---|
| Strain | 20 | 0.462 | 17.42 | 19.61 | **<0.001** |
| Parity | 1 | 0.012 | 9.36 | 4.45 | **<0.001** |
| Residuals | 397 | 0.526 | – | – | – |

**Table 8** Procrustes MANOVA testing the independent effects of strain and parity (parous vs non-parous) on female pelvic shape. Significance assessed by residualrandomization with 1,000 permutations. Type III sums of squares. Female pelvic shape differed among strains ($R^2 = 0.462$, $p = 0.001$) and showed a smallerbut significant effect of parity after accounting for strain ($R^2 = 0.012$, $p = 0.001$).

| Effect | Df | $R^2$ | F | Z | p |
|---|---|---|---|---|---|
| Strain | 20 | 0.489 | 19.40 | 15.88 | **<0.001** |
| Parity | 1 | 0.010 | 7.73 | 5.85 | **<0.001** |
| Residuals | 397 | 0.501 | – | – | – |

**Table 9** Procrustes MANOVA testing the independent effects of strain and parity on female neurocranial morphology. Significance assessed by residual randomization with 1,000 permutations. Type III sums of squares. Neurocranial morphology was primarily structured by strain ($R^2 = 0.489$, $p = 0.001$), with a modest but significant parity effect ($R^2 = 0.010$, $p = 0.001$).

| Effect | Df | $R^2$ | F | p |
|---|---|---|---|---|
| Strain | 20 | 0.404 | 12.29 | **<0.001** |
| Residuals | 362 | 0.596 | – | – |

**Table 10** Procrustes MANOVA testing the effect of strain on male pelvic shape.Significance assessed by residual randomization with 1,000 permutations. Type I sums of squares. Male pelvic shape differed strongly among strains ($R^2 = 0.404$, $p = 0.001$).

| Effect | Df | $R^2$ | F | p |
|---|---|---|---|---|
| Strain | 20 | 0.486 | 17.10 | **<0.001** |
| Residuals | 362 | 0.514 | – | – |

**Table 11** Procrustes MANOVA testing the effect of strain on male cranial shape. Significance assessed by residual randomization with 1,000 permutations. Type I

sums of squares. Male cranial shape differed strongly among strains ($R^2 = 0.486$, $p = 0.001$).

| Analysis | r-PLS | Effect Size (Z) | p-value |
|---|---|---|---|
| Pelvis-skull 2B-PLS, sex-pooled (corrected for sex, strain and parity) | 0.267 | 3.6211 | **<0.001** |
| Pelvis-skull PLS, females only (corrected for strain and parity) | 0.338 | 3.2403 | **<0.001** |
| Pelvis-skull PLS, males only (corrected for strain) | 0.226 | 0.4924 | 0.293 |

**Table 12** results of the 2B-PLS of skull shape and pelvis shape.

| Generation | r | Slope | p-value | MSE |
|---|---|---|---|---|
| F0 | 0.263 | 0.188 | **<0.001** | $5.01X10^{-5}$ |
| F1 | 0.185 | 0.101 | **0.011** | $3.17X10^{-5}$ |
| F2 | 0.406 | 0.196 | **<0.001** | $5.98X10^{-5}$ |
| B1 | 0.342 | 0.192 | **<0.001** | $4.30X10^{-5}$ |
| B2 | 0.067 | 0.030 | 0.54 | $4.35X10^{-5}$ |
| B3 | 0.327 | 0.141 | 0.12 | $2.99X10^{-5}$ |

**Table 13** correlation coefficients of skull scores and pelvis scores obtained from the 2B-PLS of skull shape and pelvis shape for both sexes.

| Generation | r | Slope | p-value | MSE |
|---|---|---|---|---|
| F0 | 0.327 | 0.162 | **<0.001** | $3.11X10^{-5}$ |
| F1 | 0.058 | 0.0217 | 0.55 | $1.96X10^{-5}$ |
| F2 | 0.268 | 0.147 | 0.37 | $9.06X10^{-5}$ |
| B1 | 0.360 | 0.161 | **<0.001** | $5.67X10^{-5}$ |
| B2 | 0.055 | 0.0215 | 0.73 | $4.29X10^{-5}$ |
| B3 | 0.0004 | 0.00022 | 0.99 | $3.73X10^{-5}$ |

**Table 14** correlation coefficients of skull scores and pelvis scores obtained from the 2B-PLS of skull shape and pelvis shape for females.

| Generation | r | Slope | p-value | MSE |
|---|---|---|---|---|

| | | | | |
|---|---|---|---|---|
| F0 | 0.179 | 0.0712 | 0.071 | 2.96X10$^{-5}$ |
| F1 | 0.240 | 0.0778 | **0.032** | 2.89X10$^{-5}$ |
| F2 | 0.378 | 0.106 | 0.052 | 2.72X10$^{-5}$ |
| B1 | 0.254 | 0.0862 | **0.005** | 4.49X10$^{-5}$ |
| B2 | 0.138 | 0.0363 | 0.37 | 3.59X10$^{-5}$ |
| B3 | 0.086 | 0.0464 | 0.81 | 1.44X10$^{-5}$ |

**Table 15** correlation coefficients of skull scores and pelvis scores obtained from the 2B-PLS of skull shape and pelvis shape for males.

| **Generation** | **Df** | **$R^2$** | **F** | **Z** | **P-value** |
|---|---|---|---|---|---|
| F0 | 1 | 0.0747 | 5.654 | 2.319 | **0.005** |
| F1 | 1 | 0.0028 | 0.274 | −0.668 | 0.728 |
| F2 | 1 | 0.0764 | 0.910 | 0.222 | 0.411 |
| B1 | 1 | 0.0310 | 4.631 | 2.352 | **0.007** |
| B2 | 1 | 0.1027 | 4.236 | 2.254 | **0.010** |
| B3 | 1 | 0.1833 | 2.469 | 1.371 | 0.103 |

**Table 16** Results of the Procrustes regression for non-parous parentals F0, hybrids F1 and F2 and backcross (B1–B3) females testing the association between pelvis Boas coordinates (shape + size) and neurocranial size. Both pelvic coordinates and neurocranial centroid size were residualized to remove effects of strainand body weight prior to analysis. Statistics are from residual randomization permutation tests (1,000permutations).

| **Predictor** | **Df** | **$R^2$** | **F** | **Z** | **p-value** |
|---|---|---|---|---|---|
| Sex | 1 | 0.08332 | 131.7585 | 6.6011 | **<0.001** |
| Strain | 20 | 0.17012 | 13.4511 | 18.5277 | **<0.001** |
| Weight | 1 | 0.00470 | 7.4355 | 3.3864 | **<0.001** |
| Age | 105 | 0.09770 | 1.4714 | 3.9838 | **<0.001** |

**Table 17** regression of Procrustes aligned pelvis coordinates on sex, weight, strain and age. All covariates had a significant effect on pelvis shape. Sex accounted for 8% of pelvis shape variance.

| **Generation** | **Procrustes Distance** |
|---|---|
| F0 | 0.05473266 |
| F1 | 0.05315449 |

| | |
|---|---|
| F2 | 0.04074632 |
| B1 | 0.04750500 |
| B2 | 0.03641539 |
| B3 | 0.02853500 |

**Table 18** Procrustes distance between mean female pelvis shape and mean male pelvis shape, after correcting for body weight, strain and age. The highest Procrustes Distance was F0, the lowest B3.

| Landmark R | Landmark L | Anatomical Definition |
|---|---|---|
| 2 | 1 | Superior point of post-tympanic hook |
| 4 | 3 | Paroccipital process |
| 6 | 5 | Posterior point on internal pterygoid process |
| 14 | 13 | Lateral point on frontal suture |
| 16 | 15 | Lateral zygomatic-frontal suture |
| 18 | 17 | Posterior zygomaticofrontal junction |
| 20 | 19 | Posterior margin of malar process |
| 22 | 21 | Frontal-temporal-parietal junction |
| 23 | 24 | Anterior margin of incisive foramen |
| 25 | 26 | Medial maxilla-premaxilla junction |
| 27 | 28 | Anterior inferior zygomatic |
| 29 | 30 | Anterior temporo-zygomatic junction |
| 31 | 32 | Anterior superior alveoli |
| 33 | 34 | Posterior incisive foramen |
| 35 | 36 | Point along palatine-maxillary suture |
| 37 | 38 | Medial palatal-pterygoid junction |
| 39 | 40 | Posterior superior alveoli |
| 41 | 42 | Lateral palatal-pterygoid junction |
| 43 | 44 | Spheno-occipital synchondrosis |
| 45 | 46 | Anterior foramen ovale |
| 47 | 48 | Posterior temporo-zygomatic junction |
| 49 | 50 | Auditory-temporal-sphenoid junction |
| 51 | 52 | Anterior inferior auditory bulla |
| 53 | 54 | Occipital-auditory-sphenoid junction |
| 55 | 56 | Point along occipitomastoid suture |
| 57 | 58 | Medial occipital condyle |
| 59 | 60 | Anterior nasal and premaxilla |
| 61 | 68 | Frontal suture on orbital rim |
| 62 | 69 | Superior temporo-zygomatic suture |
| 63 | 70 | Posterior zygomatic process |
| 64 | 71 | Superior posterior tympanic ring |
| 65 | 72 | Occipital-auditory junction |

| | | |
|---|---|---|
| 66 | 67 | Midline superior incisor |
| 91 | 79 | Posterior tympanic ring |
| 80 | 81 | Anterior inferior maxilla |
| 82 | 85 | Medial point of first upper molar |
| 83 | 86 | Medial point of second upper molar |
| 84 | 87 | Medial point of third upper molar |
| 89 | 90 | Midline |

**Table 19.** Paired cranial fixed landmarks

| Landmark | Anatomical Definition |
|---|---|
| 7 | Posterior point of presphenoid |
| 8 | Superior most point of foramen magnum |
| 9 | Posterior most point of occipital |
| 10 | Lambda |
| 11 | Bregma |
| 12 | Nasion |
| 73 | Anterior foramen magnum |
| 74 | Midline junction between basioccipital and sphenoid |
| 75 | Midline junction between sphenoid and presphenoid |
| 76 | Anterior junction of the endocranial presphenoid |
| 77 | Endocranial junction between frontal and ethmoid |
| 78 | Anterior most point of nasal bone |
| 88 | Anterior point on alveolar process between incisors |

**Table 20.** Midline cranial fixed landmarks

| Landmark | Anatomical definition |
|---|---|
| 1 | Promontorium |
| 2 | S1 Body, Posterior |
| 3 | S1 Spine |
| 4 | S2 Spine |
| 5 | Caudion, Posterior |
| 6–14 | Sacral curvature |
| 15 | Caudion, Anterior |

**Table 21.** Midline sacral landmarks

| Landmark L | Landmark R | Anatomical definition |
|---|---|---|
| 16 | 45 | Caudion, Lateral |
| 17 | 46 | Inferior Sacro-Iliac Junction |
| 18 | 47 | Superior Articular Facet: Lateral Superior |
| 19–28 | 48–57 | Alar-auricular ridge curvature |
| 29–38 | 58–67 | Lateral ridge curvature of the sacral body |
| 39–44 | 68–73 | Sacro-iliac joint curvature, posterior |
| 74 | 145 | Superior Pole, Pubic Symphysis |
| 75 | 146 | Iliospinale |
| 76 | 147 | Inferior Sacro-Iliac Junction |
| 77–91 | 148–162 | Iliac crest |
| 92 | 163 | Ischiospinale |
| 93–102 | 164–173 | Sciatic notch |
| 103 | 174 | Inferior Symphyseal Point |
| 104 | 175 | Ischiale |
| 105–113 | 176–184 | Ischial tuberosity |
| 114 | 185 | Obturator Tubercle Point |
| 115–124 | 186–195 | Foramen obturator |
| 125 | 196 | Anterior Symphyseal Point |
| 126 | 197 | Posterior Symphyseal Point |
| 127 | 198 | Iliospinale, anterior inferior |
| 128 | 199 | Pubic Eminence Point |
| 129–138 | 200–209 | Iliopubic ridge |
| 139–143 | 210–214 | Anterior superior pubic ridge |
| 144 | 215 | Superior point of the sacro-iliac joint |

**Table 22.** Left and right paired landmarks.
